# Mechanical interaction enables a collective mode of protocell proliferation

**DOI:** 10.64898/2026.04.11.717946

**Authors:** Yisen Li, Yilin Wu

## Abstract

The proliferation of primitive life-forms (known as protocells) without sophisticated cell-division molecular machineries has been an intriguing question in evolutionary biology, synthetic biology, and living matter physics. While various modes of protocell proliferation at the individual level have been proposed, the growth dynamics of protocell colonies (i.e., protocolonies) received less attention. Here we chose to study this question in protocolonies consisting of densely packed protocells derived from wall-deficient L-form bacteria. We discovered that protocolonies proliferated robustly under spatial confinement, while isolated protocells failed to divide and eventually experienced membrane rupture due to imbalance of surface and volume growth. Combining results from quantitative imaging and computational modeling, we attributed this unexpected finding to mechanical shearing between densely packed protocells driven by their growth activities; such mechanical shearing enhances cell deformation, thereby enabling cell division and sustaining population growth in a protocolony. Our study reveals a unique role of self-generated mechanical stresses in the lifestyle of primitive life-forms. The findings may help to understand and control the collective growth dynamics of synthetic protocells or active droplets.

## Introduction

Protocells refer to compartmentalized, primitive life-forms [1] that can perform metabolism and proliferate via rudimentary physicochemical pathways. Protocells are thought to have existed prior to the emergence of the last common universal ancestor (LUCA) in the evolutionary history of life [2, 3]. As protocells lack modern cell-division machineries, namely the FtsZ-Min system that controls binary fission in bacteria [4] and the spindle apparatus that drives mitosis of eukaryotic cells [5], how protocells proliferate has been an intriguing question in evolutionary biology [6], synthetic biology [7, 8] and living matter physics [9, 10].

Wall-deficient bacteria (known as L-form bacteria) perhaps best resemble protocells [11, 12], while giant lipid vesicles [3, 13] and membraneless droplets [9, 14] are also commonly used to model protocells. Experiments with both L-form bacteria and giant lipid vesicles have suggested various modes of spontaneous protocell division involving dynamic membrane deformation [6], such as extrusion-resolution [15-19] and internal vesiculation [20-22]. However, most studies on protocell proliferation based on wall-deficient bacteria were focused on the behavior at the single-cell level. Lacking motility mechanisms for dispersal, protocells would develop into densely packed aggregates (hereinafter referred to as protocolonies) during proliferation, where cells are in close proximity and could interact mechanically. The growth dynamics in such densely packed protocolonies are still unclear. Mechanical interaction between cells in a protocolony may impact each other’s membrane fluctuation and potentially modify their growth pattern, since dynamic membrane deformation is expected to play crucial roles in protocell proliferation [16-18].

Here we study the growth dynamics in colonies of bacterial protocells derived from wall-deficient *Bacillus subtilis* [15-17, 23]. Surprisingly, we found that the proliferation of bacterial protocells under spatial confinement depends on the packing density: protocolonies consisting of densely-packed wall-deficient *B. subtilis* proliferated robustly, whereas a population of isolated bacterial protocells tended to die out via membrane rupture. To understand this finding, we quantified the growth and morphological dynamics of the highly irregular, amoeba-like cells in densely packed conditions by meticulous segmentation; together with membrane tension measurement and computational modeling, our results suggest that growth-induced mechanical stresses in the protocolony promote cell-shape deformation and cell division, both of which help to maintain the surface-volume balance and sustain population growth. Our study reveals a unique role of self-generated mechanical stresses in the lifestyle of primitive life-forms. The findings may help to understand and control the collective growth dynamics of synthetic protocells [1, 7, 8] or active droplets [24, 25].

## Results

### Proliferation of bacterial protocells depends on packing density

To investigate the long-term growth dynamics of bacterial protocells derived from wall-deficient *B. subtilis* [15-17, 23], we sandwiched the bacterial protocells between the bottom surface of a multi-well culture plate and a double-layered agar pad (Fig. S1A; Methods). This culture condition confines cells in a quasi-two-dimensional space and prevents flow-induced cell drifting, thus allowing continuous single-cell tracking for up to ∼14-18 hr. We found that the prosperity of bacterial protocells depended on local cell density. As shown by the results of nucleic acid staining with SYTO16 (Methods), protocolonies consisting of densely-packed wall-deficient *B. subtilis* were able to proliferate indefinitely until running out of space (Fig. 1A, upper; Fig. 1B, red lines); also see Movie S1, left and middle panels. By contrast, well-isolated protocells tended to die out due to lysis (i.e. rupture of cell membrane and release of cytoplasmic contents) after a period of biomass growth (Fig. 1A, lower; Fig. 1B, green line; Movie S2, left and middle panels), with a lifespan of 305±140 min (mean±S.D, N=291) (Fig. S1B). This result indicates that protocolonies proliferated via a mechanism independent of the previously established proliferation mechanisms for isolated *B. subtilis* L-form cells [16, 17], since isolated cells here mostly terminated by lysis.

**Fig. 1.**
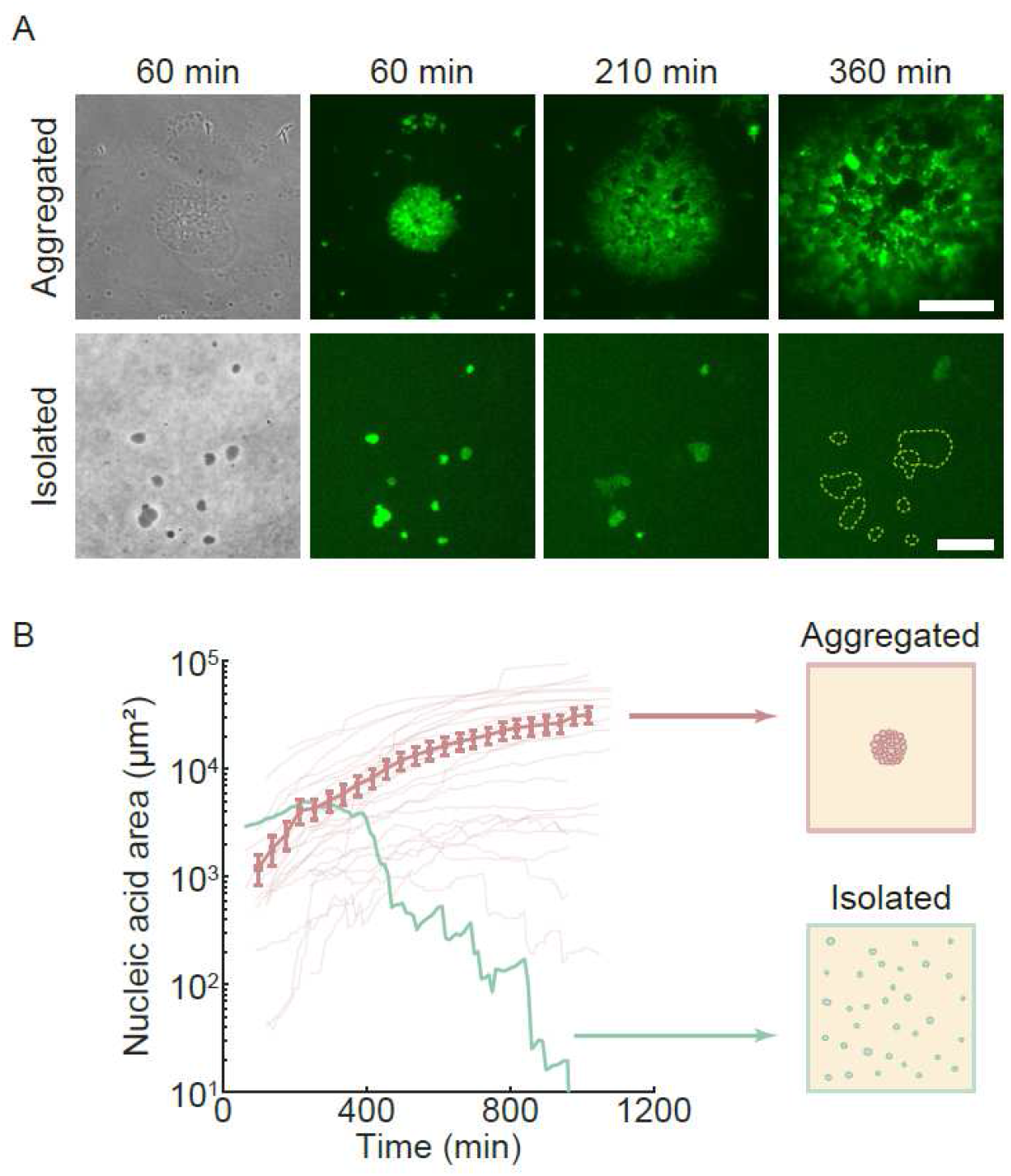
Growth dynamics of bacterial protocells in densely packed aggregates and in isolation. (A) Image sequences showing the development of bacterial protocells derived from wall-deficient *B. subtilis* in an aggregated protocolony (upper) and in isolation (lower). The data was from representative cases. The first panel in each row is a phase-contrast image. The green false color displays the fluorescence from nucleic acids (stained by SYTO16; Methods), which served as a proxy of the biomass of cells. For cells in protocolonies, the area occupied by nucleic acids kept increasing, while scattered hollow areas appeared within the protocolony due to cell death; also see Movie S1. For isolated protocells, the area occupied by nucleic acids was decreasing and cells gradually lysed; lysis events were identified by sudden loss of membrane integrity (Movie S2; Methods). Note that the area with weak fluorescence enclosed by dashed lines are nucleic acids remnants of recently lysed cells; the fluorescence disappeared shortly as nucleic acids diffused away. Scale bars: 50 μm (upper panels) and 20 μm (lower panels). (B) Temporal dynamics of nucleic acid area during the growth of cells in protocolonies (lines and data points in red color) and of isolation protocells (green line). The nucleic acid area was computed from SYTO16 fluorescence images (Methods) and it served as a proxy of biomass. For protocolonies (N=27), each light-colored line represents data from one protocolony; the data points connected by straight lines are the averaged biomass of all protocolonies, with the error bars representing the standard error of the mean. For isolated protocells, the sum of nucleic acid area in a population (N=291) was plotted. The schematic diagrams illustrate the spatial distribution of aggregated and isolated cells. In both panels T=0 min corresponds to the time of inoculation.

To understand this surprising result, we sought to compare the growth dynamics of individual cells in protocolonies and of isolated protocells. However, performing cell segmentation in protocolonies for the single-cell measurement in long-term time-lapse imaging was challenging, because cells had highly irregular amoeba-like shapes and the boundary between cells was hardly distinguishable from their cytoplasm either under phase-contrast microscopy or in fluorescence imaging of commonly used lipophilic membrane stains (such as FM 4-64 [26]). We serendipitously found that the oxidative stress sensor CellROX Deep Red labelled cell cytoplasm and displayed brighter fluorescence at the interface between neighboring bacterial protocells (Movie S1, right panel; Methods); moreover, its spectrum can be paired with that of the nucleic acid stain SYTO16 for multichannel imaging. With the help of this fluorescent dye and a machine learning algorithm for pattern recognition (Methods), we were able to perform efficient segmentation of densely packed protocells.

Using the cell segmentation data and single-cell tracking, we measured the growth dynamics of individual cells and fitted the volume growth curves into an exponential form ∼exp(*βt*), where *β* is defined as the volume growth rate (Methods). We found that the average volume growth rate of cells in protocolonies (0.50 ± 0.67 *h*^−1^, mean±S.D., N=1762) was not significantly different from that of isolated protocells prior to lysis (0.38 ± 0.32 *h*^−1^,mean±S.D., N=291) (Fig. S2A), despite that the volume growth rate distribution of cells in protocolonies displayed a longer tail (i.e., a portion of cells had much higher growth rates than others in the population) (Fig. S2B,C). We further examined the single-cell growth dynamics by categorizing the cells into different growth patterns based on a combination of growth-curve fitting parameters. Counterintuitively, isolated cell populations had a greater proportion undergoing active growth than cells in protocolonies (64% vs 49%), whereas 37% of cells in protocolonies remained constant in size (versus 22% for isolated cells) (Fig. S3). These results raised a question: why did protocells proliferate better in densely packed environments than in isolation even though isolated cells grew more actively?

### Aggregated protocells are capable of maintaining surface-volume balance

Since the lifespan of well-isolated bacterial protocells is mostly terminated by cell lysis, understanding the mechanism of cell lysis may provide clues to the origin of the difference in proliferation capability between the two growth modes of bacterial protocells. As programmed cell death in bacteria is generally induced by environmental stresses, which are not present in our growth condition, we reasoned that cell lysis occurred when the speed of surface area increase could not keep up with that of the volume increase. To examine this idea, we quantified the morphological dynamics of bacterial protocells. We measured the surface area increase speed (*dS*/*dt*) and the volume increase speed (*dV*/*dt*) of single cells during protocell growth, and computed their ratio relative to a perfect circular disk of the same volume that underwent the same volume growth (denoted as the surface-volume balance ratio *η* (*t*); Methods). A surface-volume balance ratio *η* < 1 corresponds to the situation that the current value of *dS*/*dt* is not sufficient to keep up with the cell’s volume increase speed; by contrast, for *η* > 1, greater values of *η* indicate that the current value of *dS*/*dt* is more likely to maintain the cell’s current shape. We found that as isolated cells grew, their average *η* decreased from 1.11±0.07 (mean±S.D., N=593) to 1.01±0.08 (mean±S.D., N=580), approaching the critical value *η*=1 (Fig. 2A, green). This result indicates that the ability of isolated cells to keep surface-volume balance declined over time. On the other hand, the *η* of aggregated cells in protocolonies during active growth only declined slightly from 1.13±0.07 (mean±S.D., N=1476) to 1.10±0.12 (mean±S.D., N=1441) (Fig. 2A, red), which is consistent with the experimentally observed low cell lysis rate in protocolonies. Thus, our result supported that imbalance of surface and volume growth underlies the higher likelihood of cell lysis of isolated cells.

**Fig. 2.**
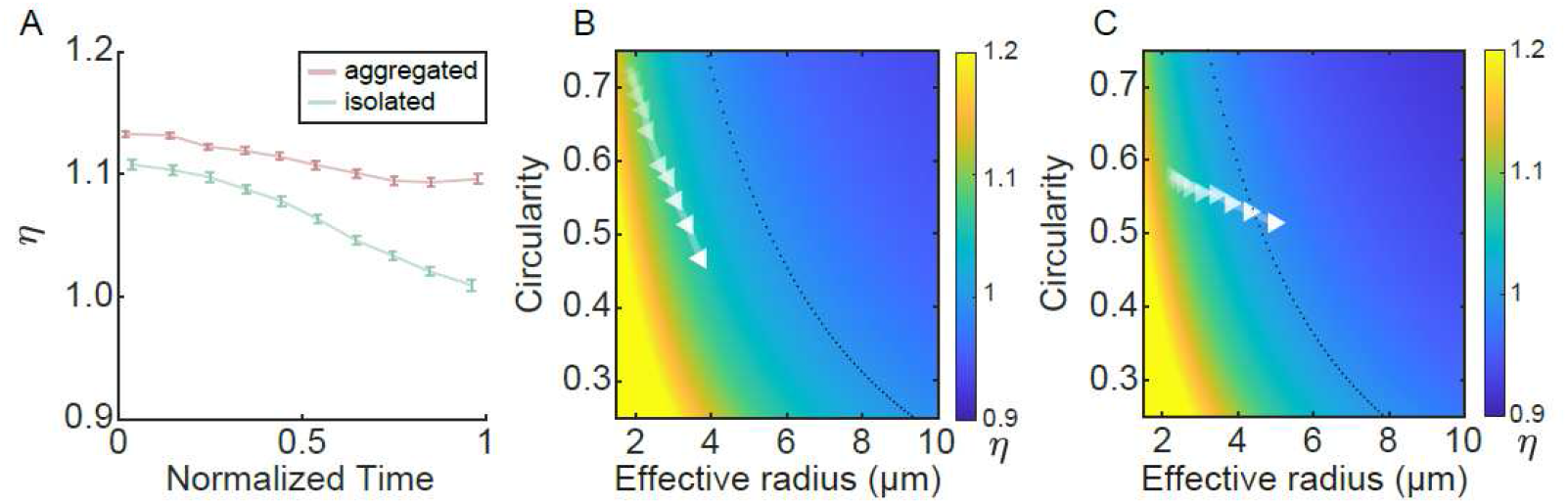
Analysis of surface-volume balance for actively growing bacterial protocells. (A) The surface-volume balance ratio (*η*) plotted against growth time for actively growing cells (see Fig. S3, panels C,H). Red and green points correspond to cells in protocolonies and isolated protocells, respectively. The time is normalized by the maximal observation period (>4 hr), or by the lifespan of a cell if it lysed within the observation period. Lines connecting the data points serve as guides to the eyes. Error bars represent standard error of the mean (N= 866 and 187 cells in protocolonies and in isolation, respectively). (B,C) Dependence of *η* on geometrical parameters (colormap) and the evolutionary trajectory of growing bacterial protocells in the geometrical space (white triangles). Panel B and C correspond to cells in protocolonies and in isolation, respectively. Colormap in each panel was computed based on our mathematical analysis and the experimentally measured growth parameters (averaged growth rates of surface area and volume) (Methods); it shows the dependence of *η* on effective radius and circularity, with the color bar to the right of each panel indicating the value of *η* and the black dotted line indicating the contour of *η=*1. The white triangles in each panel represents the average effective radius and circularity of cells at different times of growth as shown in panel A (time increasing from left to right).

More generally, mathematical analysis shows that the value of *η* for a pancake-shaped cell with area *A* and perimeter *P* is controlled by two geometrical parameters, the effective radius 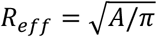 and the circularity *C* = 4*πA*/*P*^2^ (i.e., a measure of cell morphology compared to a perfect circular disk) (Methods). Using experimentally derived parameters, we plotted *η* on the plane of *R*_*eff*_ and *C* for actively growing cells in protocolonies (Fig. 2B) and for isolated protocells (Fig. 2C), respectively. As shown in Fig. 2B,C, the average circularity *C* of both cells in protocolonies and isolated protocells decreased during volume growth (indicated by the increase of *R*_*eff*_). However, the trajectory of *η* for cells in protocolonies remained above the critical value *η*=1 (dashed line in Fig. 2B), while the trajectory of *η* for isolated cells reached below *η*=1 (dashed line in Fig. 2C). To conclude, the analysis suggests that cells in protocolonies are on average more capable of maintaining surface-volume balance and thereby reducing the probability of cell lysis.

### Growing protocells deform and divide due to mechanical interaction

In addition to preventing lysis, bacterial protocells have to divide in order to sustain the growth of a population. While division was seldom observed in isolated cells, cells in protocolonies were able to divide at the later stage of growth (> ∼2.5 hr after inoculation) (Fig. S4) and consequently, they had a significantly smaller size than the isolated counterparts (Fig. 3A; Fig. S5). We note that the average circularity of cells in protocolonies decreased steadily over time, and the variance of the circularity distribution for cells in protocolonies was much higher than that for isolated cells over the entire time course of growth (Fig. 3B), indicating that a portion of cells in protocolonies had a low circularity (i.e., a large extent of cell shape deformation) (Fig. 3B). Interestingly, data obtained with cells in protocolonies show that the division rate and the extent of cell shape deformation were positively correlated (Fig. 3C). Since the division of bacterial protocells is independent of the FtsZ-related mechanism [15] and it requires the formation of lipid membrane stalks (the fusion site of two lipid bilayer membranes when they approach each other prior to scission) [27], we attributed this correlation to the fact that cell shape deformation increases the likelihood of membrane stalk formation.

**Fig. 3.**
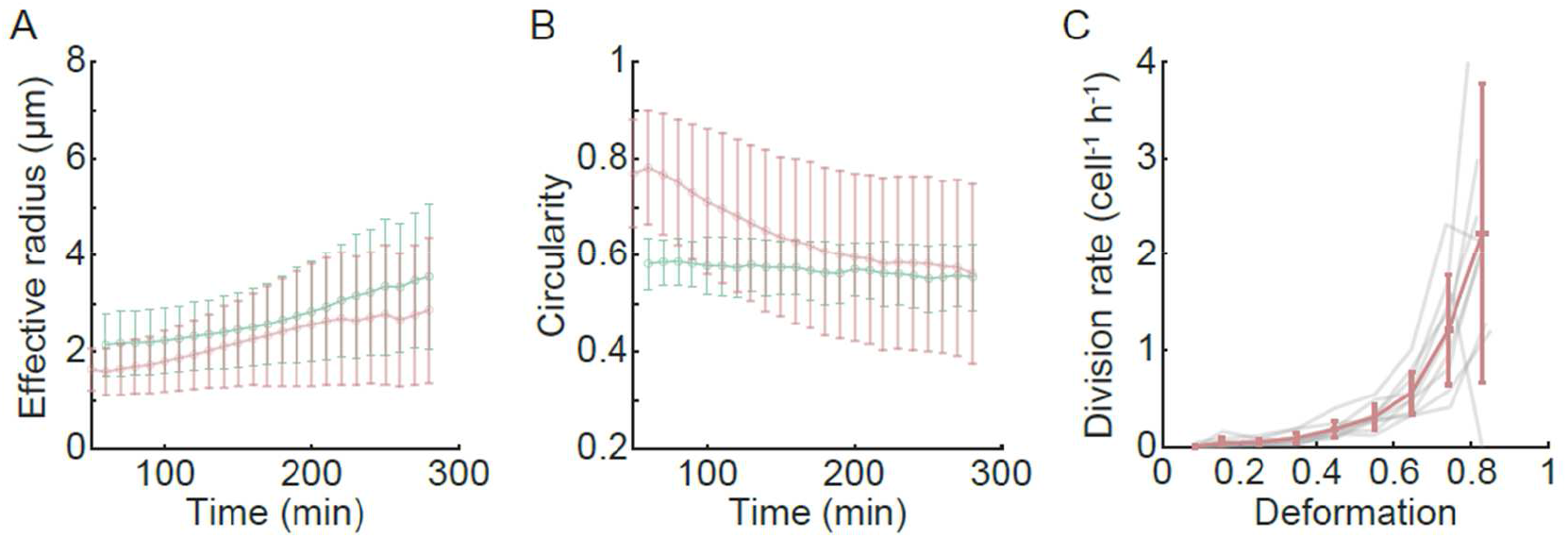
Correlation between shape deformation and division of bacterial protocells. (A,B) Effective radius (panel A) and circularity (panel B) of cells plotted against the growth time of bacterial protocells. Red and green data correspond to cells in protocolonies and isolated cells, respectively. T=0 min corresponds to the time of inoculation. Error bars represent standard deviation (N>50 cells). Lines connecting the data points serve as guides to the eyes. For data in panel A, two-tailed Mann-Whitney-Wilcoxon test was performed to examine whether each pair of red and green data sets at a specific time point were drawn from the same distribution, and the test showed that every pair of data sets were significantly different (*P* < 0.05). For data in panel B, Levene’s test was performed to examine the equality of variances of each pair of red and green data sets at a specific time point, and the test showed that the variances of every pair of data sets were significantly different (*P* < 0.05). Also see Fig. S5. (C) Protocell division rate (number of divisions per cell per hour) plotted against the extent of cell deformation defined as (1 – C), where C represents the circularity of cells. Data were acquired in 9 protocolonies (gray lines); red line shows the average of data from all gray lines, with the error bars indicating standard deviation. For each protocolony, all cells and all the division events throughout the entire observation period were included in the analysis.

The sub-population of cells that experienced large deformation (decreasing circularity) and were able to divide may be a key driver of population growth in protocolonies. We sought to understand how this subpopulation arose in the protocolony. In general, cell shape deformation is driven by spatiotemporal variation of tension on the cell membrane. So we measured the membrane tension for cells in protocolonies using a fluorescent probe FliptR via Fluorescence Lifetime Imaging Microscopy (FLIM) (Methods). When inserted into lipid membranes, the structure of FliptR may be altered by membrane tension variation; consequently, the fluorescence lifetime of FliptR is positively correlated with membrane tension [28]. For cells in protocolonies, the FliptR fluorescence lifetime measurement revealed that the membrane tension of their lateral (or peripheral) surface in contact with others was higher than that of the rest part of the cell membrane (referred to as the bulk part), and both were significantly higher than that of isolated protocells (Fig. 4). Given the indistinguishable average volume growth rates between cells in protocolonies and isolated protocells (Fig. S2), the higher membrane tension of cells in protocolonies is presumably because the cells are mechanically sheared by actively growing neighbors in the crowded environment; such mechanical shearing would increase membrane tension primarily at the interface between cells (i.e., the peripheral surface) but weakly in the bulk part of the membrane due to substrate pinning. This growth-induced mechanical shearing between cells may enhance the likelihood of large deformation and enables cell division.

**Fig. 4.**
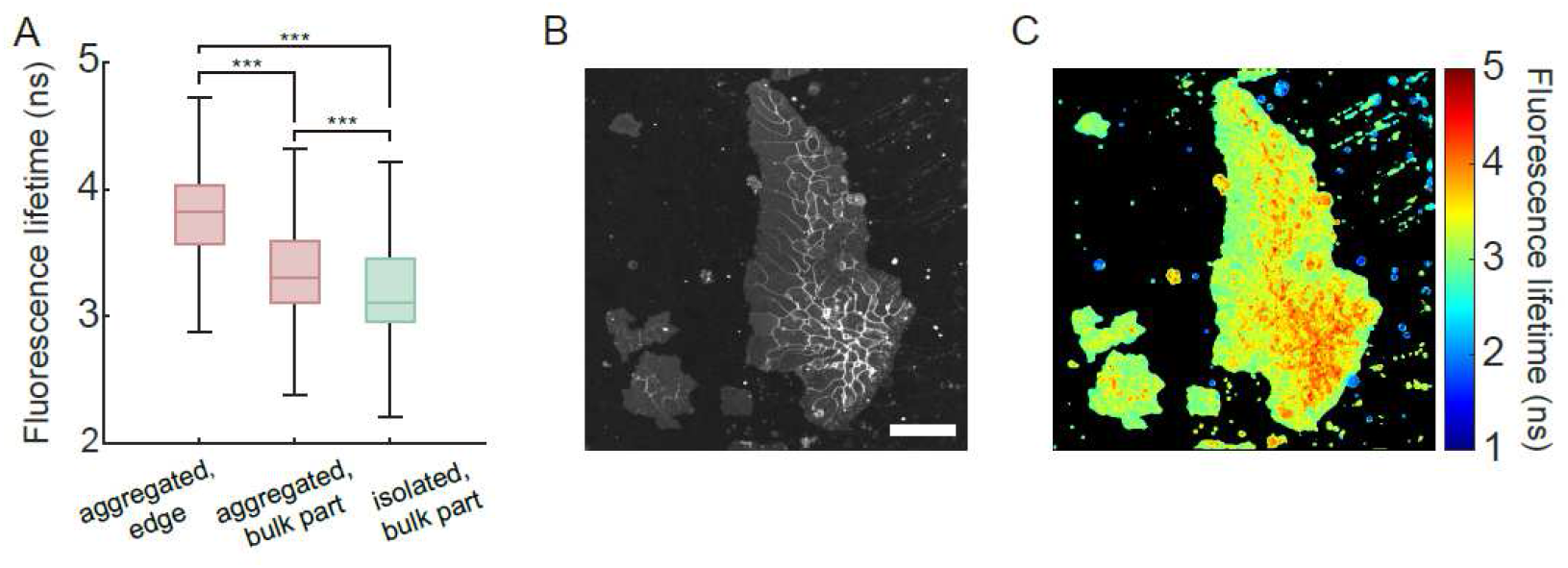
Membrane tension measurement based on the fluorescent probe FliptR. (A) Boxplot showing the fluorescence lifetime distributions of FliptR on bacterial protocell membranes. Due to the quasi-2D interstitial confinement (Fig. 1A), most of a cell’s membrane is in contact with the double-layered agar pad and the multi-well culture plate; this part of cell membrane is denoted as “bulk part”. The membrane at the interface between neighboring cells in a protocolony (i.e., brighter lines in the fluorescence intensity plot in panel B) is denoted as “edge”. Fluorescence lifetime of FliptR on the membrane was analyzed pixel-wise for 5 protocolonies and 162 isolated cells (Methods), and data from all the pixels were grouped and plotted here. The line inside each box represents the median, the top and bottom edges of each box represent the quartiles, and the ends of whiskers represent the largest (or lowest) data point within 1.5-fold of the interquartile range (i.e., the box height) from the upper (or lower) edge of the box. Significance test was performed by two-tailed Mann-Whitney-Wilcoxon non-parametric test for the equality of means of data sets that do not follow Gaussian distributions (\*\*\**P* < 1.0 × 10^−10^). (B) Fluorescence intensity image of FliptR staining cell membrane in protocolonies. The higher fluorescence intensity at the interface between neighboring cells than the bulk part of cells is due to the higher density of lipid membrane at cell-cell contact areas. Scale bar, 20 μm. (C) Spatial distribution of FliptR fluorescence lifetime obtained by FLIM imaging for the protocolonies shown in panel B. False color scale represents fluorescence lifetime in units of nanoseconds.

### Computational modeling supports the mechanical origin of collective protocell proliferation

The experiments above suggest that mechanical shearing causes large deformation in a subpopulation of protocolonies, thereby enabling cell division and sustaining population growth. To examine this notion, and to further understand the developmental dynamics of protocolonies, we developed a Cellular Potts Model [29, 30] to simulate the growth dynamics of protocolonies in quasi-2D environment (Fig. 5A,B). Here bacterial protocells were modeled as active droplets whose dynamics evolve according to the Monte Carlo algorithm, which involves a free energy function controlling the interactions, growth, and morphology of protocells; in addition, the surface-volume balance ratio *η* controls the cell death rate (Methods). We made two modifications to the commonly used free energy function in Cellular Potts Model (Methods): first, to prevent cell volume collapse during cell-cell compression, we adopted an osmotic pressure contribution [31] instead of the commonly used quadratic effective energy in the free energy function; second, to account for the experimental observation that protocells are able to squeeze through densely packed neighbors by means of large deformation, we introduced a history-dependent substrate-pinning energy.

**Fig. 5.**
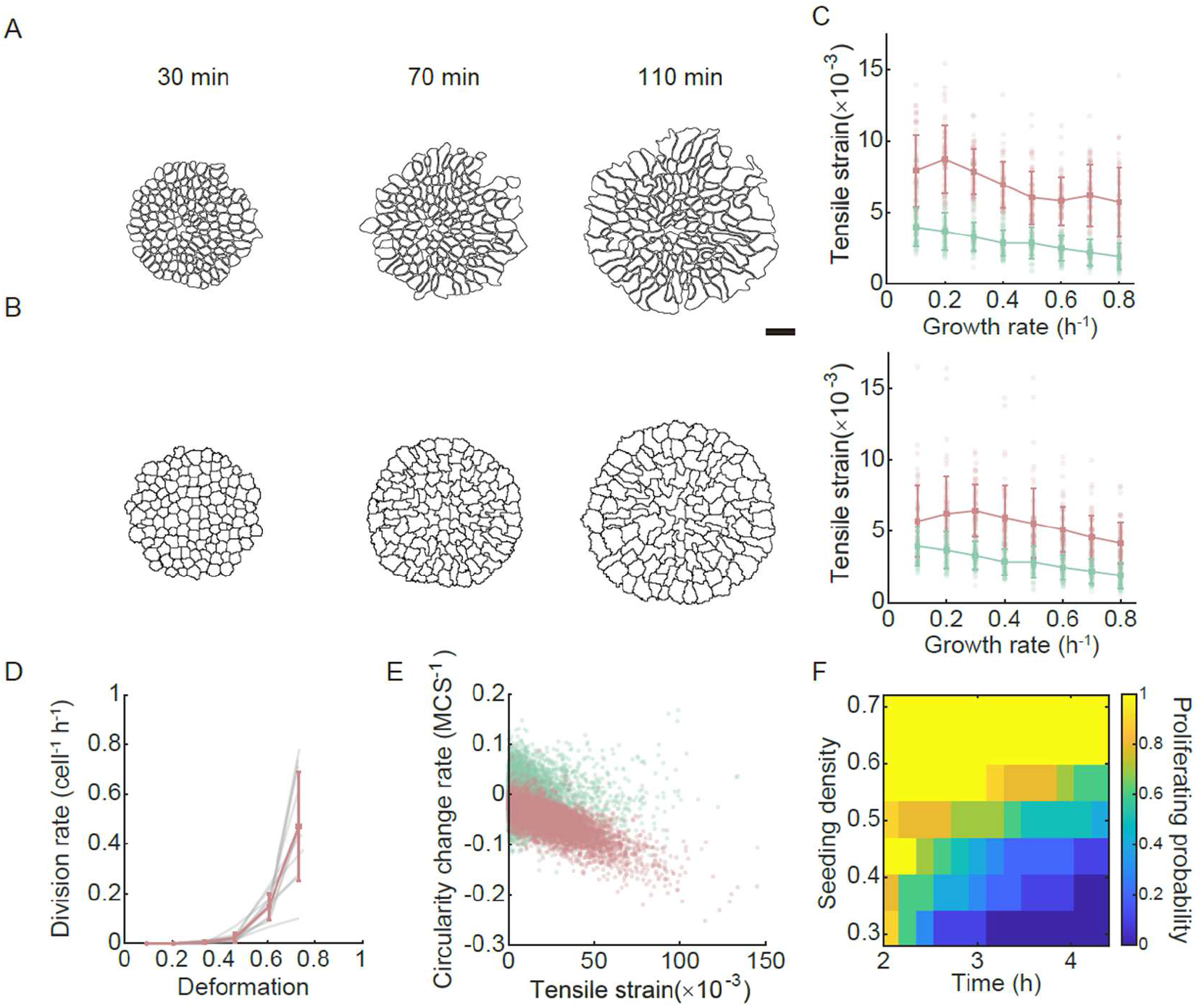
Cellular Potts Model simulations of protocell growth dynamics. (A,B) Comparison of protocolony morphology between experiment (panel A) and Cellular Potts Model simulation (panel B). The experimental image sequence shown in panel A was segmented according to the fluorescence images of CellROX-stained cells (Methods). To initiate the simulation 120 circular cells were seeded at hexagonal close-packing density (Methods). Scale bar, 10 μm. (C) Membrane tensile strain of protocells plotted against growth rate in simulations. Red and green data correspond to cells in protocolonies and in isolation, respectively. For cells in protocolonies, the membrane tensile strain is computed for the periphery area (upper panel) and the bulk part (lower panel); for isolated protocells, the overall membrane tensile strain is computed (see Methods). Scattered points represent the time-averaged membrane tensile strain and the prescribed growth rate of individual cells. Data points in the line plots represent the mean membrane tensile strain of cells with the same prescribed growth rate, with error bars representing standard deviation of membrane tensile strain (N > 45 simulated cells). Lines connecting the data points serve as guides to the eyes. (D) Simulated protocell division rate (number of divisions per cell per hour) plotted against the extent of cell deformation defined as (1 – C), where C represents the circularity of cells. Data were acquired in 11 protocell populations (gray lines); red line shows the average of data from all gray lines, with the error bars indicating standard deviation. The analysis for each protocell population was performed by taking into account all cells and division events across all simulation steps, following the same calculation method as in Fig. 3C. (E) Rate of circularity change plotted against membrane tensile strain in simulations. Red and green data correspond to cells in protocolonies and in isolation, respectively. Each scattered point represents the instantaneous membrane tensile strain (calculated in the same manner as in the upper panel of Fig. 5C) and the instantaneous rate of circularity change for a cell. The Spearman correlation coefficient of membrane tensile strain and circularity change rate is -0.69 (p ≈ 0) and -0.02 (p=0.04) for cells in protocolonies and in isolation, respectively. (F) Time evolution of protocell proliferating probability depends on seeding density. The seeding density was varied by tuning the average cell-cell distance in simulated protocell populations, with a density of 1.0 corresponding to hexagonal close packing (Methods).

We calculated tensile strain on the membrane of simulated cells (Methods) and used it as a measure of membrane tension. Over a wide range of cell mass growth rates, we found that cells in protocolonies generally experience higher membrane tensile strain than those of isolated cells at the same growth rate (Fig. 5C); in particular, the peripheral membrane tensile strain (Fig. 5C, upper) is greater than that of the bulk part (Fig. 5C, lower). These results are in agreement with the experimental data in Fig. 4A. Furthermore, our simulations reproduce the positive correlation between the extent of cell shape deformation and the cell division rate (Fig. 5D, as compared to the experimental data in Fig. 3C). We find that a faster decrease of cell circularity (i.e., a greater rate of cell shape deformation) in protocolonies is associated with higher membrane tensile strain (Fig. 5E). These simulation results support the notion that shearing in protocolonies can substantially increase membrane tension and promote cell division by deforming cells.

As the extent of cell shearing due to external mechanical stress depends on local cell density, we expect that the seeding density of protocells at the time of inoculation would affect the long-term proliferation of the entire population. The model allows us to continuously vary the initial seeding density of protocells, which was not attainable in our experiments (see Methods). Indeed, our simulations show that protocell populations proliferate more robustly at higher seeding density (Fig. 5F): The proliferating probability (defined as the surviving proportion in an ensemble of simulated protocell populations; Methods) of protocell populations remains ∼1.0 over time for seeding densities above ∼60% of hexagonal close-packing, but it gradually diminishes to zero at lower seeding densities. In the latter case, the lifetime of the entire protocell population increases with the seeding density.

## Discussion

In this study, using protocells derived from wall-deficient bacteria, we discovered that self-generated mechanical stress enables a collective mode of protocell proliferation. Compared to their isolated counterparts, cells in protocolonies consisiting of densely-packed prococells were capable of maintaining the surface-volume growth balance and more importantly, they were able to divide; these two factors reduce the likelihood of cell lysis and allow the protocolony to proliferate in a collective manner. Membrane tension measurement and Cellular Potts modeling suggest that cell division in protocolonies is facilitated by mechanical shearing between cells in the crowded environment due to their growth activities. Taken together, our results reveal a unique role of self-generated mechanical stresses in the proliferation of protocells.

The collective mode of protocell proliferation enabled by self-generated mechanical stresses may be relevant to the propagation of L-form bacteria derived from densely packed pathogens [32] escaping from wall-targeting antibiotic treatment [33] and phage infection [34], such as *Enterococci* biofilm infections in urinary tract and wounds [35, 36]. This mechanism of collective L-form proliferation is independent of two previously established proliferation mechanisms for isolated L-form cells, namely lipid membrane overproduction [17] and increased membrane fluidity [16], since isolated cells mostly terminated by lysis in the quasi-two-dimensional growth environment. Presumably, the substrate pinning in confined space suppresses spontaneous membrane deformation arising from either lipid membrane overproduction or increased membrane fluidity. Note that Gram-negative bacteria L-forms have two layers of lipid membrane, with the outer membrane being a rigid load-bearing shell [19, 37, 38]; the self-generated mechanical stress due to growth activity may be insufficient to deform the double-layered membrane in Gram-negative L-forms, and consequently could not proliferate by the mechanism we report here.

More generally, our study may shed light on the lifestyle of primitive life forms or protocells that existed prior to the emergence of modern cell-division machineries [2, 3]. Protocells are believed to thrive in porous natural structures in early Earth environment, such as porous rocks in hydrothermal vents [39, 40] and mica sheets [41], because such structures could enrich minerals and organic molecules that facilitated essential biochemical reactions or served as building blocks for protocells [42]. In these hypothetical hotspots of primitive life forms, the confined space would limit protocell dispersal and give rise to densely packed protocolonies. The collective proliferation mechanism enabled by self-generated mechanical stresses might have been a viable means of protocell propagation in such restricted physical environments, where cells were in close proximity and interacted mechanically.

A long-standing goal in origins-of-life research is to generate artificial cells that can autonomously grow and divide [3, 7, 8]. Achieving this goal would not only help to elucidate the fundamental principles of life but will also provide a synthetic biology platform for biomedical applications [43]. While recent development in synthetic biology has proposed various strategies to generate the full cell cycle by bottom-up approaches [7, 8], devising the functional molecular machinery for cell division is still a daunting task. Instead, the collective proliferation mechanism we report here enables cell division that requires neither dedicated division machineries nor external perturbation, such as temperature variation [44], external mechanical shear [18], or nonequilibrium chemical reactions [45]. Therefore, it offers a simple solution to achieve spontaneous propagation of synthetic cells before engineered division machinery and cell size regulation modules become available.

## Methods

### Bacterial culture and protocell preparation

To produce stable wall-deficient (L-form) bacterial cells as a model of protocells, we used *B. subtilis* LR2 carrying a xylose-inducible murE expression cassette where the native *murE* promoter was replaced by a xylose-inducible promoter [15] (gift from J. Errington, Newcastle University). This strain was maintained in the form of walled-cells in Luria-Bertani (LB) medium (1% Bacto tryptone, 0.5% yeast extract, and 0.5% NaCl) supplemented with D-(+)-Xylose (0.5% wt./vol.) and chloramphenicol (5 μg/mL). To synchronize the transition from walled-state to wall-deficient cells, *B. subtilis* LR2 walled-cells were artificially turned into protoplasts (i.e., cells with the peptidoglycan cell wall stripped off) by lysozyme treatment following a previously published protocol [46]; when the protoplasts resume growth and can propagate stably in wall-deficient state in the absence of xylose, they are used as model protocells in this study. Specifically, single-colony isolates of *B. subtilis* LR2 walled-cells from 1.5% wt./vol. Bacto agar plates were inoculated in 3 mL LB supplemented with D-(+)-Xylose (0.5% wt./vol.) and grown overnight with gyration at 180 rpm at 30 °C to stationary phase. 60 μL of the overnight culture was diluted in 3 mL of the xylose-supplemented LB medium and further incubated with shaking for 3-4 hours, yielding an exponential-phase culture with an optical density (at 600 nm) of ∼0.2. Walled cells in the exponential-phase culture were harvested and re-suspended in 3 mL xylose-free L-form medium (see below) supplemented with lysozyme (1 mg/ml) and incubated with shaking at 37 °C for 2-3 h. The L-form medium was prepared by mixing 2-fold-concentrated (2×) MSM medium (40 mM MgCl_2_, 1 M sucrose and 40 mM maleic acid, adjusted to pH 7.0 with 10 M NaOH; denoted as 2×MSM) with 2× LB broth (i.e., 2% Bacto tryptone, 1.0% yeast extract, and 1.0% NaCl) at a 1:1 ratio. The L-form medium (denoted as “LB/MSM”) was an osmoprotective nutrient medium modified from the Nutrient Broth/MSM medium [15]. The obtained protoplasts in 1 mL LB/MSM were centrifuged at ∼350 × g (2000 r.p.m of an Eppendorf benchtop centrifuge) for 10 min and resuspended gently in 50-200 μL fresh LB/MSM for further experiments. In all subsequent experiments xylose was not supplemented to the growth media such that cells remained in wall-deficient state.

### Preparation of double-layered agar pad for bacterial protocell growth

The double-layered agar pad consists of a thick layer of 4.0% Bacto agar and a thin layer of 0.6% Bacto agar, both of which are infused with LB/MSM described above. The 0.6% Bacto agar layer was sandwiched between the 4.0% Bacto agar layer and a solid substrate; in addition to promoting cell growth, the thin 0.6% LB/MSM agar layer reduces the likelihood of sliding of the 4.0% LB/MSM agar layer during cell growth, thus allowing long-term observation of cell growth. To prepare the double-layered agar pad, we first melted solidified 8.0% wt./vol. Bacto agar infused with 2×LB medium in a microwave oven. The molten agar (∼70 °C and ∼60 °C for 4% and 0.6% desired agar concentrations, respectively) was mixed with prewarmed 2×MSM medium at a 1:1 ratio, yielding 4% or 0.6% LB/MSM agar. At this point, fluorescence dyes may be supplemented to the liquefied agar: to quantify cell mass and to perform cell profiling, the nucleic acid stain SYTO16 (Thermo Fisher Scientific, Cat. No: S7578) and the cytoplasm stain CellROX™ Deep Red Reagent (Thermo Fisher Scientific, Cat. No: C10422) were added into the agar at final concentrations of 0.3 μM and 1 μM, respectively; for FLIM imaging, FliptR (Spirochrome, Cat. No: SC020) was added at a final concentration of 1 μM. Then 1 mL of the 4% LB/MSM agar was pipetted onto a 24 mm × 24 mm coverslip and covered by another coverslip of the same size; the assembly of agar and coverslip was cooled for 10 min at room temperature inside a Petri-dish of 150 mm diameter with the lid on. After agar solidification, the coverslip on the top of the 4% LB/MSM agar was removed. The agar was dried under laminar airflow for 3-5 min and trimmed to a size of 15 mm × 15 mm. Next, 10 μL of the 0.6% LB/MSM agar was pipetted on top of the trimmed 4% LB/MSM agar pad and covered by a new coverslip to form a thin agar layer on top of the 4% LB/MSM agar layer. The entire assembly (double-layered agar sandwiched between two coverslips) was cooled under laminar airflow for 15 min. After the 0.6% LB/MSM agar solidified, the coverslip covering it was removed, and the entire assembly (with the 0.6% LB/MSM agar exposed to air) was further dried under laminar airflow for 1-3 min. The double-layered agar pad was finally trimmed to 7 mm × 9 mm, which was used within 5-10 min.

As soon as the double-layered agar pad was ready to use, we inoculated wall-deficient *B. subtilis* cell in the interstitial space between the double-layered agar pad and the surface of a culture well. Specifically, we deposited a total volume of 0.5 μL protoplast culture prepared above on several different spots of a rectangular culture well (9.4 mm ×10.7 mm) of a multi-well plate (µ-Slide 8 Well, ibiTreat; Cat. No: 80826) and covered the well by the prepared double-layered agar pad. When the double-layered agar pad was applied to the deposited protoplast suspension, air bubbles often appeared and cell aggregates of various sizes tended to form at the boundary of the bubbles. The dissolution of air in the bubbles (normally within 1 hr) left an inhomogeneous distribution of protoplasts, with some well isolated from others and some in densely-packed aggregates; the latter were taken as protocolonies. Similarly to the preparation of agar pad, the following fluorescence dyes may be supplemented to the prepared protoplast culture before deposition onto the culture well: to quantify cell mass and to perform cell profiling, the nucleic acid stain SYTO16 and the cytoplasm stain CellROX™ Deep Red Reagent were added into the culture at final concentrations of 0.3 μM and 1 μM, respectively; for FLIM imaging, FliptR was added at final concentrations of 1 μM.

### Phase contrast and epifluorescence imaging

Imaging was performed on motorized inverted microscopes (Nikon TI2-E). The following objectives were used in different experiments for phase contrast and epifluorescence imaging: Nikon CFI Plan Fluor DLL 40×, N.A. 0.75, W.D. 0.66 mm; Nikon Plan Apo λ 60× oil, N.A. 1.4, W.D. 0.13 mm. Note that for regions occupied by large protocell colonies, the 40× objective was used to take the large-field images by stitching overlapping images, which allows the entire protocolony to be included in the field of view. Fluorescence imaging was performed in epifluorescence using filter sets specified below, with the excitation light provided by a solid-state light engine (Lumencor SPECTRA X or Lumencor SPECTRA III). Recordings were made with an sCMOS camera (Andor Neo 5.5, Andor Technology) using the software NIS-Elements AR (Nikon). For all experiments requiring long-term protocell growth, the multi-well plates were covered with lids and sealed with Parafilm to prevent evaporation. The temperature was maintained at 30 °C unless otherwise stated, using a custom-built temperature-control system installed on the microscope stage and a lens warmer on the objectives. The lens warmer was controlled by a temperature controller (CU-501, Live Cell Instrument).

### Fluorescence lifetime measurement with laser scanning confocal microscopy

Fluorescence lifetime imaging microscopy (FLIM) of FliptR-stained bacterial protocells was performed on a laser scanning confocal microscope (Leica DMi8 inverted microscope with Leica TCS SP8 X system) via a 100× oil objective (Leica No. 506210, N.A. 1.4, W.D. 0.09 mm). Recordings were made by a hybrid detector (Leica HyD) and specific software and hardware for single molecule detection (PicoQuan). The excitation light (480 nm) was provided by a pulsed white light laser at 20 MHz. The hybrid detector was used to collect emission light at 570 nm-630 nm.

To measure the membrane tension of cells in protocolonies and in isolation, bacterial protocells and the double-layered agar pads supplemented with 1 μM FliptR were prepared following the procedures described above. The bacterial protocells were inoculated in the interstitial space between the double-layered agar pad and the bottom surface of the multi-well plate. After incubation for 6-8 hours at 30 °C, the multi-well plate was transferred to the stage of the confocal microscope and maintained at room temperature. The fluorescence images were acquired for selected regions of interest (ROIs) through the 100× oil objective. The acquisition procedures were controlled by the software platform LAS X (Leica).

### Growth dynamics of bacterial protocells

To investigate the growth dynamics of cells in protocolonies and in isolation, *B. subtilis* protoplasts were inoculated in the interstitial space between a double-layered agar pad and a culture well of a multi-well plate following the procedures as described above. The agar pad was supplemented with the nucleic acid stain SYTO16 and the cytoplasm stain CellROX Deep Red to quantify the proliferable biomass and to profile cells (for cell size calculation), respectively. SYTO16 has superior brightness and requires a relative low excitation light intensity, thus conferring photostability and low phototoxicity in long-term time-lapse imaging.

CellROX Deep Red is an oxidative stress sensor; we found that it stains cell cytoplasm and displays bright fluorescence at the interface between cells (Movie S1, right panel; Fig. S4A), allowing efficient segmentation of cells (especially in densely-packed environments). The multi-well plate was then transferred to the microscope stage (Nikon TI2-E) and cells were observed through the 40× or 60× phase contrast objectives at 30 °C. We started acquisition of microscopy images at ∼1-4 h after inoculation of cells, when the cells had adapted to the interstitial environment and resumed growth (checked by eyes under phase contrast microscopy). Microscopy images were acquired for more than 10 hr in 10-min or 15-min intervals. During each interval, a sequence of image acquisition operations was performed over ∼10 ROIs (2048 pixel × 2048 pixel). At the beginning of each time interval, the motorized stage moved to each selected ROI successively; at each ROI, we acquired a phase-contrast image, a fluorescence image of SYTO16 and a fluorescence image of CellROX Deep Red. The image acquisition procedures were performed automatically (controlled by software NIS-Elements AR; Nikon) and for each interval the image acquisition for all ROIs was completed within ∼2 min. For the acquisition of fluorescence images, the excitation light was provided by a solid state light engine (Lumencor SPECTRA X or Lumencor SPECTRA III), and the following filter sets were used: a ratiometric pHluorin2 filter set for SYTO16 (excitation: 485/20 nm, FF02-485/20-25, Semrock; emission: 535/22 nm, FF01-535/22-25, Semrock; dichroic: 506 nm, FF506-Di03-25x36, Semrock) and a Cy5 filter set for CellROX Deep Red (excitation: 640/30 nm, emission: 690/50 nm, dichroic: 660 nm; 49009 - ET - Cy5 Narrow Excitation, Chroma); the excitation light was triggered by a TTL signal sent from a custom-programmed Arduino microcontroller, which received signals from a custom-written macro in NIS-Elements AR (Nikon) that controlled the entire image acquisition protocol. For the acquisition of phase contrast images, the transmitted light was provided by a built-in white light LED controlled by NIS Elements directly. All recordings were made by the Andor Neo sCMOS camera; the camera was turned on 10 ms after the fluorescence excitation light was triggered on, and the excitation light was turned off right after the camera exposure.

### Image processing and data analysis for experiments

Images were processed using the open-source Fiji (ImageJ) software (http://fiji.sc/Fiji), the open-source ilastik software (https://www.ilastik.org) [47], and custom-written programs in MATLAB (The MathWorks; Natick, Massachusetts, United States).

We quantified biomass by calculating the area occupied by SYTO16-stained nucleic acids in bacterial protocells. First, we need to identify the area of SYTO16 fluorescence. To do so we binarized the fluorescence images of SYTO16 following the steps below: (1) Align the time-lapse images by the built-in image registration methods (primarily the Registration Estimator app) in MATLAB. (2) Duplicate the raw fluorescence images in ImageJ and apply a Gaussian filter to these images (denoting the resulted images as the low frequency images). (3) Divide the raw fluorescence images by the low frequency images to reduce non-uniformity and enhance image quality (denoting the resulted images as enhanced images). (4) Import the enhanced images to ilastik, a machine-learning-based image analysis tool.

Manually label a small portion of regions as nucleic acid or background based on the fluorescence intensity as training dataset. The software would automatically identify the unlabeled regions based on the intensity and texture features of the labeled regions, segmenting nucleic acid from the background for all the enhanced images. (5) Threshold the gray scale images obtained in Step 4 and export from ilastik the binarized images with regions only labeled as nucleic acid. The identified area of SYTO16 fluorescence is taken as biomass.

The temporal evolution of biomass was analyzed as follows, after identifying the area of SYTO16 fluorescence. For cells in protocolonies (Fig. 1C, red lines), the binarized fluorescence images of SYTO16 were analyzed in the following steps: (1) Define the protocolonies in binarized SYTO16 fluorescence images by density-based spatial clustering of applications with noise (DBSCAN), a built-in function in MATLAB. The parameter epsilon is set to 45 pixels (approximating 7.5 μm), and the minimum number of neighbors within epsilon radius is 3. (2) Link the protocolonies between frames according to their overlapping nucleic acid area. Specifically, the intersection over union (IoU) of nucleic acid area from protocolonies in two consecutive frames is calculated. The IoU threshold is set to 0.2, above which two protocolonies would be linked. (3) Plot the nucleic acid area of each protocolony against time and smooth the time-area curves by the built-in filloutliers function in MATLAB. For the analysis of biomass of isolated protocells (Fig. 1C, green line), as the growth medium was supplemented with both SYTO16 and CellROX Deep Red, we first profiled the cells using the fluorescence images of CellROX-stained cell cytoplasm (see the paragraph below for cell size calculation); then we imported the binarized fluorescence images of both SYTO16 and CellROX Deep Red into the ImageJ TrackMate plugin as two separate channels. The TrackMate plugin then simultaneously tracked the temporal evolution of cytoplasm-occupied area of individual cells using the intersection over union (IoU) algorithm [48] (with each cell identified as a 4-connected region in TrackMate; see the paragraph below for volume growth dynamics analysis) using data in the CellROX Deep Red channel and computed the nucleic acid area of these cells using data in the SYTO16 channel. The tracking data were imported to MATLAB with a custom-written program so that the total nucleic acid area from a population of isolated protocells was plotted against time. The lifespan of isolated protocells (Fig. S1B) was defined as the duration from inoculation to the time of lysis; it was calculated based on the time evolution of nucleic acid occupied area of isolated cells obtained by the binarization and tracking procedures described above.

To profile bacterial protocells for cell size calculation, we made use of the CellROX Deep Red fluorescence images. For isolated cells, we binarized the fluorescence images of CellROX Deep Red following the same procedures for identifying the area of SYTO16 stained nucleic acids described above. Regions occupied by previously lysed cells were labeled as background due to the low fluorescence intensity in the training dataset for ilastik. The output data from ilastik were manually checked to correct errors. For cells in protocolonies, the CellROX fluorescence intensity was much higher at the interface between neighboring cells than in the cytoplasm (Fig. S4A); therefore, the fluorescence images of CellROX were aligned in MATLAB and manually segmented in Fiji (ImageJ) simply based on the fluorescence intensity.

To analyze the volume growth dynamics of bacterial protocells (Fig. S3), we noted that the cell volume is linearly proportional to the cytoplasm-occupied area because of the pancake shape of cells under quasi-2D confinement. The cytoplasm-occupied area was measured by cell profiling based on fluorescence images of the cytoplasm stain CellROX Deep Red. Specifically, the binarized fluorescence images of CellROX Deep Red obtained above were tracked in the ImageJ TrackMate plugin using the intersection over union (IoU) algorithm [48]; during tracking, each cell in each frame was identified as a 4-connected region in TrackMate, and the area of this region approximates the area occupied by cell cytoplasm. We imported the tracking data generated by TrackMate into MATLAB and analyzed it by custom-written programs. Using the least-squares method in MATLAB, we fitted the cytoplasm-occupied area (*A*) of each individual cell into an exponential function *A* = *A*_0_exp(*βt*), where *β* is taken as the cell volume growth rate and *A*_0_ is the initial cytoplasm-occupied area of the cell. Then we classified the growth curves of all cells (both in protocolonies and in isolation) based on the following three parameters: (1) the standard deviation of cell area in the time sequence, denoted as *SD*; (2) volume growth rate obtained by the exponential fit of the growth curve, *β*; and (3) goodness of growth rate fitting *R*^2^. Five growth patterns were defined according to the values of these 3 parameters (Fig. S3A,B): “active growth” (*SD* > 3, *β* > 0, *R*^2^ > 0.8), “modest growth” (*SD* > 3, *β* > 0, *R*^2^ ≤ 0.8), “constant” (*SD* ≤ 3), “decline” (*SD* > 3, *β* ≤ 0, *R*^2^ > 0.8), and “modest decline” (*SD* > 3, *β* ≤ 0, *R*^2^ < 0.8). For statistical analysis of the volume growth rates, the growth rate of cells following the “constant” or “modest decline” growth patterns was set to zero, since cells in the two groups barely displayed any volume change (Fig. S3F,G,J,K).

To process FLIM data, we fitted the fluorescent decay curves in Fiji (ImageJ) FLIMJ plugin [49] with Levenberg-Marquardt (LMA), two-exponential reconvolution algorithms. The instrument response function was reconstructed by SymPhoTime software in the PicoQuan imaging system and exported for use in the reconvolution in FLIMJ. The bulk part and the edge of cells were identified by binarizing the FliptR fluorescence intensity images in the software ilastk using procedures mentioned above. The binarized bulk part and the edge images were applied as masks to FLIM images obtained by FLIMJ to remove the background.

### Analysis of surface-volume balance during protocell growth

In our experiment, the bacterial protocell grew in the quasi-2D interstitial space between the double-layered agar pad and the multi-well culture plate. Taking a cell as a “pancake” with area *A*(*t*) and thickness *h*, the surface area of a cell can be approximated by *S*(*t*) = 2*A*(*t*) + *P*(*t*) · *h* and the cell volume *V*(*t*) = *A*(*t*) · *h*, where *P*(*t*) is the perimeter of the cell area *A*(*t*) and it can be calculated from the cell profiling results described above. Both *S*(*t*) and *V*(*t*) are assumed to follow exponential growth dynamics, i.e., *S*(*t*) = *S*_0_ exp(*α* t) and *V*(*t*) = *V*_0_ exp(*βt*), where *S*_0_ and *V*_0_ are the initial cell’s surface area and volume, and *α* and *β* are the surface area growth rate and volume growth rate, respectively; note that the molecular machinery for phospholipid synthesis is primarily located on the cell membrane [50], so the speed of cell membrane synthesis is expected to be proportional to cell surface area, i.e., *dS*(*t*)/*dt* ∝ *S*(*t*), resulting in the exponential growth dynamics. We introduced two geometrical parameters, the circularity 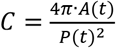 and the effective radius 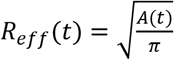, and substituted *A*(*t*) and *P*(*t*) with the two parameters in the expressions of *S*(*t*) and *V*(*t*). Thus, we have the ratio between *dS*(*t*)/*dt* and *dV*(*t*)/*dt* for protocell growth (denoted as *η*_1_(*t*)):

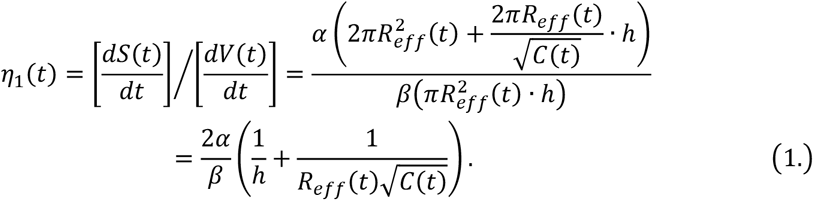

For a perfect circular disk of the same volume (i.e., with radius *R*_*eff*_(*t*) and thickness *h*) and undergoing the same volume growth dynamics, the ratio between its surface area growth speed and volume growth speed *η*_0_(*t*) is:

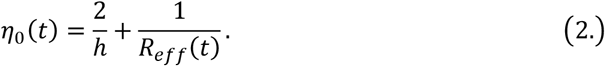

We defined the surface-volume balance ratio *η* (*t*) as follows:

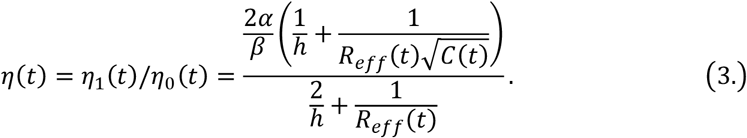

Higher values of *η* indicate that cells are more capable of keeping surface-volume balance; in particular, *η* < 1 indicates that the current surface area growth speed is not sufficient to keep up with the cell’s volume increase, even if the cell is approaching a circular disk shape that would require minimal surface area growth. To compute the *η* curves in Fig. 2A, we calculated the growth rates of surface area and volume of individual cells (*α* and *β*) by fitting the experimentally obtained temporal dynamics of *S*(*t*) and *V*(*t*) into exponential functions (using the least-squares method in MATLAB) in the form of *S*(*t*) = *S*_0_ exp(*αt*) and *V*(*t*) = *V*_0_ exp(*βt*); we also calculated their *R*_*eff*_(*t*) and *C*(*t*) based on the single-cell area tracking data obtained by the procedures described above. The cell thickness *h* was chosen to be 0.8 µm, equivalent to the width of walled cells. The colormaps of *η* in Fig. 2B and Fig. 2C were obtained with Eq. 3 (substituting the single-cell growth rates of surface area and volume *α* and *β* in the equation with their ensemble average) for cells in protocolonies and for isolated protocells, respectively.

### Cellular Potts Model simulation and analysis

The Cellular Potts Model is a lattice-based framework for modeling cell dynamics that preserves cell-cell interactions and deformable cell shape [29, 30]. We modeled cells as growing grid patches in a square lattice with periodic boundary conditions. Each modeled cell with cell index *σ* (or “spin”) initially occupied a circular area of *A*_*σ*_ (i.e., the initial cell size); the spin of the environment (i.e., non-occupied lattice sites) was set as *σ* = 0. Each lattice site assumes a “type” *τ* that denotes either the modeled cell (*τ* = 1) or the environment (*τ* = 0). Cells can grow and move by randomly choosing a lattice site on the cell boundary and copying its spin into one of its 4-connected neighboring lattice sites. The spin-copying attempt, also known as the “spin-flipping” attempt, was accepted with a probability related to difference between the global energies before and after the attempt (denoted as ℋ_before_ and ℋ_after_, respectively), following the Monte Carlo Metropolis algorithm [51],

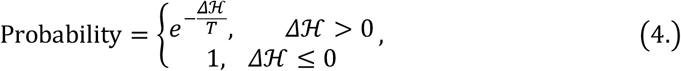

Where Δℋ = ℋ_after_ − ℋ_before_ and T is an effective temperature. A Monte Carlo step (MCS) contains numerous spin-copying attempts (Table S1). The definition of global energy ℋ and the criteria for determining cell division and lysis are described below.

The global energy ℋ in our model consists of boundary energy ℋ_Boundary_, area free energy ℋ_Area_, perimeter free energy ℋ_Perimeter_ and a substrate-pining energy ℋ _Substrate_:

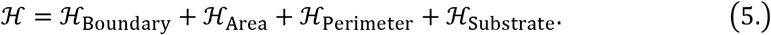

The boundary energy is described as follows:

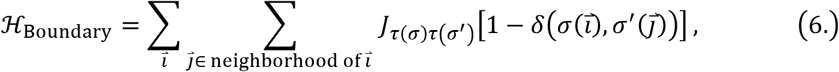

where *J* is the contact energy (see Table S1), (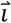is a pixel position of spin *σ*,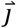 is the Moore neighborhood (8-connected neighborhood) of (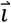, and *τ* (*σ*) is the type of spin *σ*. The *δ* is the Kronecker delta function:

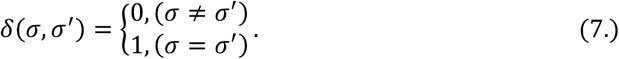

Note that *J* is set to be inversely related to the distance between neighboring lattices, in order to reduce lattice anisotropy due to the unequal distance of lattice sites in the 8-connected neighborhood in the square lattice. Experimentally, a cell cannot lose its area without cell-cell interactions unless lysis happens; therefore, the modeled cell occupied pixel (*τ* = 1) was set to reject the replacement by the environment (*τ* = 0).

The area free energy was used to calculate the penalty for deviations of the current area from the target area; here the area corresponds to the volume of cells in 3D but not the membrane area. The function of ℋ_*Area*_ is adapted from the reference [31]:

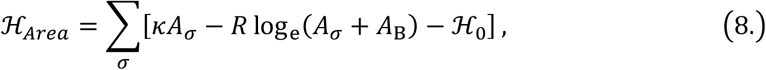

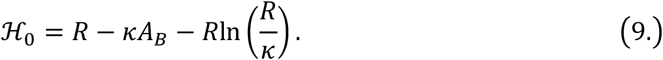

In Eq. 8 *A*_*σ*_ is the current area of cell *σ* (with *σ* > 0) and *A*_*B*_ is a buffering area. *K* and *R* are parameters that control the shape of the function, and ℋ_0_ is a value that shift the minimum of this function. Eq. 8 is different from the commonly used quadratic effective energy to describe volume conservation; it includes an osmotic pressure contribution [31] that prevents cell volume collapse during cell compression.

Nonetheless, denoting the target area of cell *σ* as 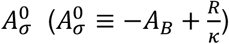, which can be found by minimizing the function [*κA*_*σ*_− *R* log_e_(*A*_*σ*_+ *A*_*B*_)], one can expand ℋ_*Area*_ near 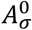 and obtain an approximate expression for ℋ_*Area*_ to recover a quadratic effective energy:

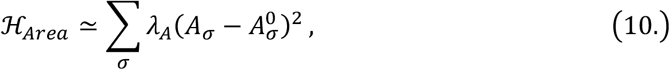

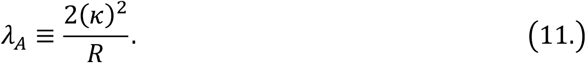

Here *λ*_*A*_ is referred to as the weight of volume growth. The target area follows exponential growth as described below:

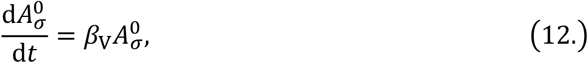

where *β*_V_ is the cell volume growth rate. Note that Eq. 8 but not Eq. 10 was used for the calculation of the area free energy.

The perimeter free energy ℋ_*Perimeter*_ was used to calculate the penalty for deviations of the current cell perimeter *P*_*σ*_ from the target perimeter 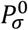:

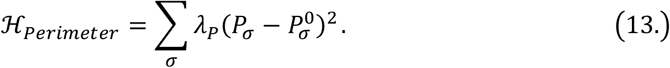

Here *λ*_*p*_ is the weight of perimeter growth. The perimeter energy prevents large cell distortion and fragmentation. The target perimeter was calculated recursively following the difference equation below:

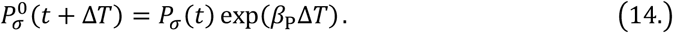

Here *P*_*σ*_ (*t*) is the perimeter of cell *σ* in the current MCS (at time *t*); 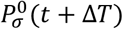 is the target perimeter of cell *σ* at the next MCS (at *t* + *ΔT*); *β*_*P*_ is the perimeter growth rate, and its value was set to be the same as that of volume growth rate *β*_*V*_; and *ΔT* is the time duration corresponding to each MCS and it can be directly compared to the time in experiments. Note that the number of attempts per MCS as specified in Table S1 was set to ensure that the volume increase precisely follows the exponential growth in Eq. 12.

The history dependent substrate-pinning energy ℋ_Substrate_ is introduced to model the substrate pinning of cell boundary in interstitial space that suppresses fusion due to spontaneous membrane deformation at the boundary of cells and yet allows large deformation when a cell is forced to deform during interaction with neighbors. The substrate-pinning effect is treated as an energy barrier contributing to total energy difference before and after a spin-flipping attempt. Each lattice site is assigned a weight of substrate-pinning energy, denoted as *L*. For the two lattice sites involved in a spin-copying or spin-flipping attempt (i.e., a lattice site on the cell boundary and one of its 4-connected neighboring lattice sites), denoting their positions as 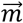 and 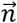, respectively, the change of the substrate-pinning energy is

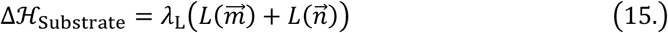

where *λ*_*L*_ is constant coefficient. *L* is set as zero during initialization; for a lattice site at position 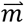 that becomes part of cell boundary or has belonged to the cell boundary in the past, its weight of substrate-pinning energy 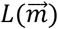 evolves according to the following equation:

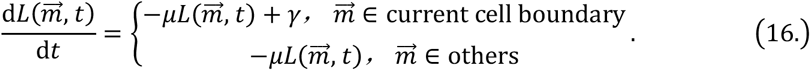

Here *µ* and *γ* are constants. Eq. 16 describes increase of substrate-pinning energy at a constant rate as soon as a lattice site becomes part of the current boundary; it also assumes exponential decay of the substrate-pinning effect. The difference equation of Eq. 16 is used to solve for 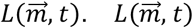 is updated only at the beginning of every MCS, so time interval of integrating Eq. 16 is *Δ*T, the time duration corresponding to each MCS. In experiments, a growing cell can push through the middle of two adjacent and connected cells, by forcing them to separate from each other. To account for this experimental observation, we set the coefficient in substrate-pinning energy *λ*_*L*_ to be 80% lower at lattice sites whose Moore neighborhood (8-connected neighborhood) contains spins that belong to two or more other cells (corresponding to tri-cellular or higher-order vertices).

The parameters used in the simulations are listed in Table S1. We calibrated the cell volume growth rate *β*_V_ in Eq. 12 in our model to ensure that one Monte Carlo Step (MCS) of cell growth corresponds to 5 minutes in the experiment. To initialize the simulations, cells were seeded randomly in the simulation domain, with both the initial cell size *A*_*σ*_ and the initial distance between cells following Gaussian distributions (Table S1). The simulation domain was a 500×500 square lattice with periodic boundary conditions; the domain size was sufficiently large to ensure that the modeled cells remained far away from the boundary throughout the entire simulation.

Cell lysis was introduced after 20 MCS based on the surface-volume balance ratio *η* (*t*) (defined in Eq. 3 above) measured for each cell. Specifically, for an individual cell, if it kept *η* (*t*) < 1 for five consecutive MCS after 20 MCS, the type *τ* of lattice sites occupied by this cell would be set to *τ* = 0 (i.e., the type of environment), which corresponds to cell lysis.

Cell division was detected when a modeled cell consisted of two or more disconnected regions at the end of each MCS. The region with the largest area was taken as the mother cell, and thus it retained the index or spin value of the mother cell; the remaining regions were regarded as newly divided daughter cells and assigned new index values.

The membrane tensile strain (Fig. 5C,E) was calculated at the end of each MCS. The membrane tensile strain of a cell was proxied by the difference between its current membrane area and its predicted membrane area at the current time. Assuming that the cells have a “pancake” geometry as in experiments, the membrane area of the *σ*-th cell, denoted as *S*_*σ*_, consists of two bases of area *A*_*σ*_ and a lateral surface of height *h*. Thus, the total membrane area is:

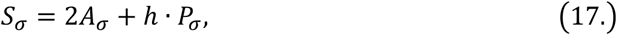

where *P*_*σ*_ is the perimeter of the *σ*-th cell. The predicted membrane area 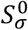 is:

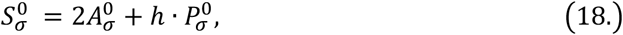

where the 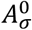 and 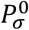 are the target area and target perimeter that evolves according to Eq. 12 and Eq. 14, respectively. For isolated cells, we compared *S*_*σ*_ and 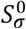 to estimate the overall membrane tensile strain (Fig. 5C,E, green data points), which is defined as follows:

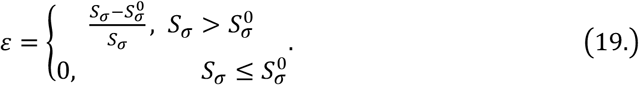

For cells in protocolonies, the membrane tensile strain is computed for the periphery area and the bulk part separately by replacing 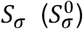 in Eq. 19 with 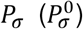 or 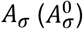, respectively. For simplicity, in all simulations for the membrane tensile strain calculation, we turned off cell lysis.

To study the effect of seeding density of cells on the long-term proliferation of protocell populations, we seeded cells with hexagonal packing within a circular region. By calculating the area occupied by the hexagonally packed cells in a triangular unit, we have the packing fraction 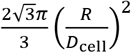, where *D*_cell_ is the mean of initial distance between cells measured from cell centers and *R* is the average seeding cell radius. The seeding density was varied by changing the initial distance *D*_cell_. The inoculum size (i.e., the number of seeded cells) was chosen as 20. The proliferating probability at a specific time is defined as the surviving proportion in an ensemble of simulated protocell populations.

## Data, Materials, and Software Availability

All study data are included in the article and/or supporting information. The custom codes used in this study are available from the corresponding author upon request.

## Acknowledgements

We thank Jeff Errington and Ling Juan Wu at Newcastle University, Romain Mercier at Institut de Microbiologie de la Méditerranée Aix-Marseille Université-CNRS, Masaki Osawa at Duke University, and Akiko Kashiwagi at Hirosaki University for their kind gifts of bacterial strains and instructions on L-form culture. This work was supported by the National Natural Science Foundation of China (NSFC no. T2425007 to Y.W.), the Ministry of Science and Technology of China (no. 2021YFA0910700, to Y.W.), and the Research Grants Council of Hong Kong SAR (RGC ref. nos. 14309023, 14307822, 14307821, RFS2021-4S04 and CUHK Direct Grants, to Y.W.). Y.W. acknowledges support from New Cornerstone Science Foundation through the Xplorer Prize.

## Author Contributions

Y.L. designed the study, performed experiments, analyzed and interpreted the data, and developed the computational model. Y.W. conceived and designed the study, analyzed and interpreted the data. Y.W. wrote the paper with input from Y.L.

## Supplementary Information (SI) for

### SI Figures

**Fig. S1.**
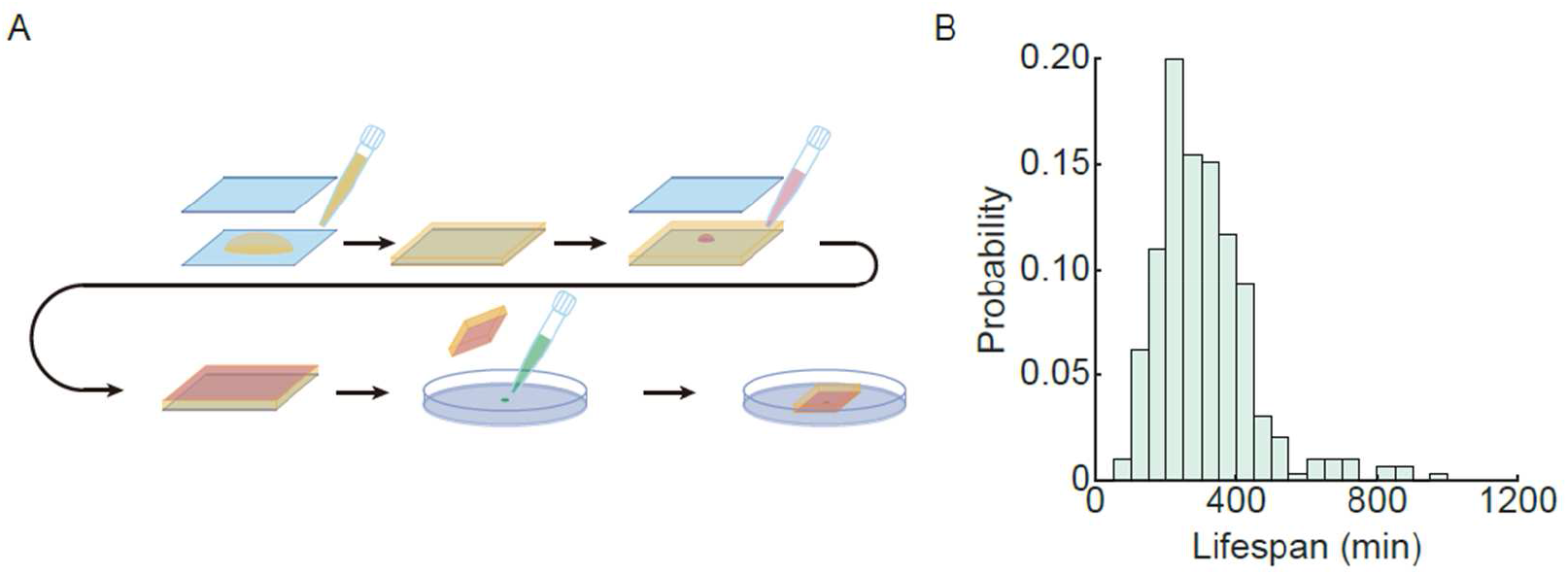
Experimental setup for long-term observation of bacterial protocell growth. (A) Schematic diagram of the quasi-2D culture method of bacterial protocells derived from wall-deficient *B. subtilis* L-forms. The double-layered agar pad consists of a thick layer of 4.0% agar (yellow) and a thin layer of 0.6% agar (red), both of which are infused with an osmoprotective nutrient medium (Methods). The thick layer of 4% agar (yellow) was solidified between two coverslips (blue). The thin layer of 0.6% agar (red) was solidified between the 4% agar and a coverslip. The prepared double-layered agar was then placed on top of the protocell suspension (green) deposited on a multi-well culture plate. The thin 0.6% agar layer, with a thickness of ∼5 µm, is in direct contact with cells and the multi-well culture plate’s bottom surface; it reduces sliding of the 4% agar pad and allows long-term observation of cells. (B) Lifespan distribution of isolated bacterial protocells. The lifespan of an isolated protocell is defined as the duration from inoculation to lysis.

**Fig. S2.**
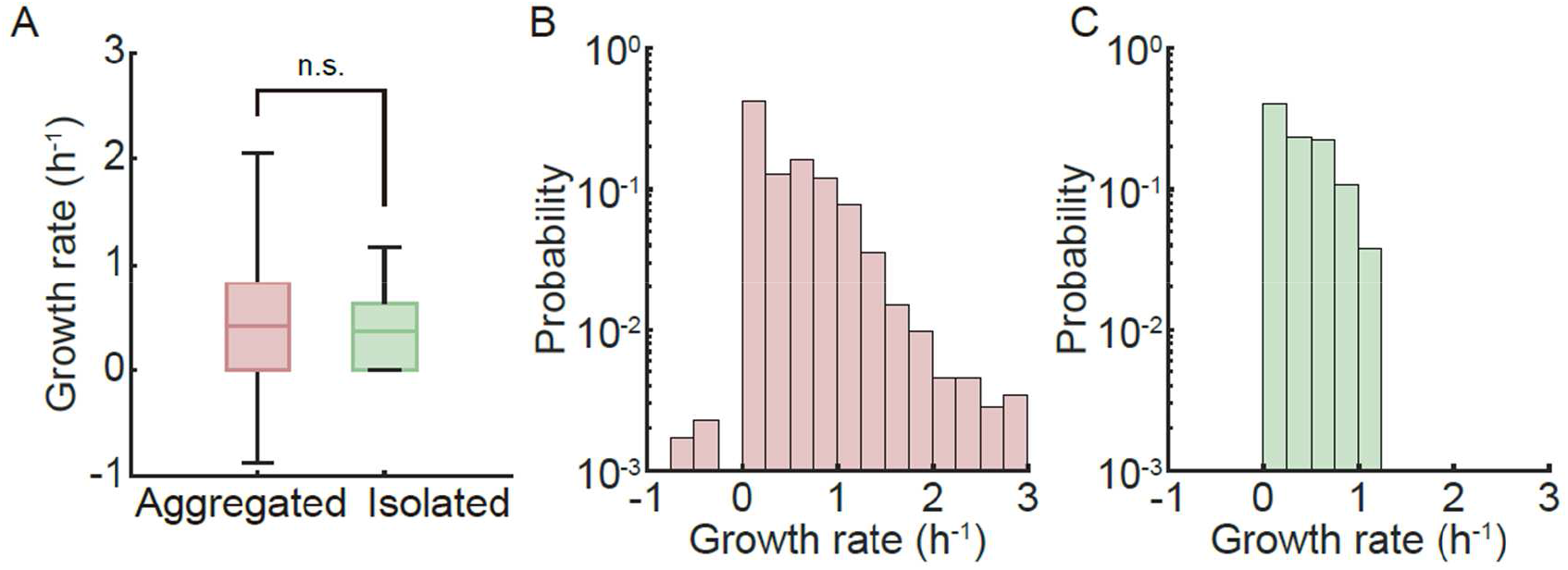
Growth rate of bacterial protocells in densely packed aggregates and in isolation. (A) Boxplot showing the volume growth rate of cells in aggregated protocolonies (red) and of isolated protocells (green). The cytoplasm-occupied area (*A*) of each individual cell is proportional to cell volume because of the pancake shape of cells under quasi-2D confinement. The cytoplasm-occupied area was measured by cell profiling based on fluorescence images of the cytoplasm stain CellROX Deep Red (Methods). It was fitted exponentially as *A* = *A*_0_exp(*βt*), where *β* is defined as the volume growth rate and *A*_0_ is the cell’s cytoplasm-occupied area at the beginning of observation (Methods; Fig. S4). The average volume growth rate of cells in protocolonies and in isolation was 0.50 ± 0.67 *h*^−1^ (mean±S.D., N=1762) and 0.38 ± 0.32 *h*^−1^ (mean±S.D., N=291), respectively. Statistical features in the boxplot are interpreted in the same way as in Fig. 4A. Significance test was performed by two-tailed Mann-Whitney-Wilcoxon test for data sets following non-Gaussian distribution. n.s., not significant. (B,C) Distribution of volume growth rate *β* of cells in protocolonies (panel B) and in isolation (panel C). A negative growth rate here indicates a reduction in cell size, which was likely due to cell mass loss caused by shearing of neighboring cells. The statistical features of the distributions are shown in panel A. Also see Fig. S3.

**Fig. S3.**
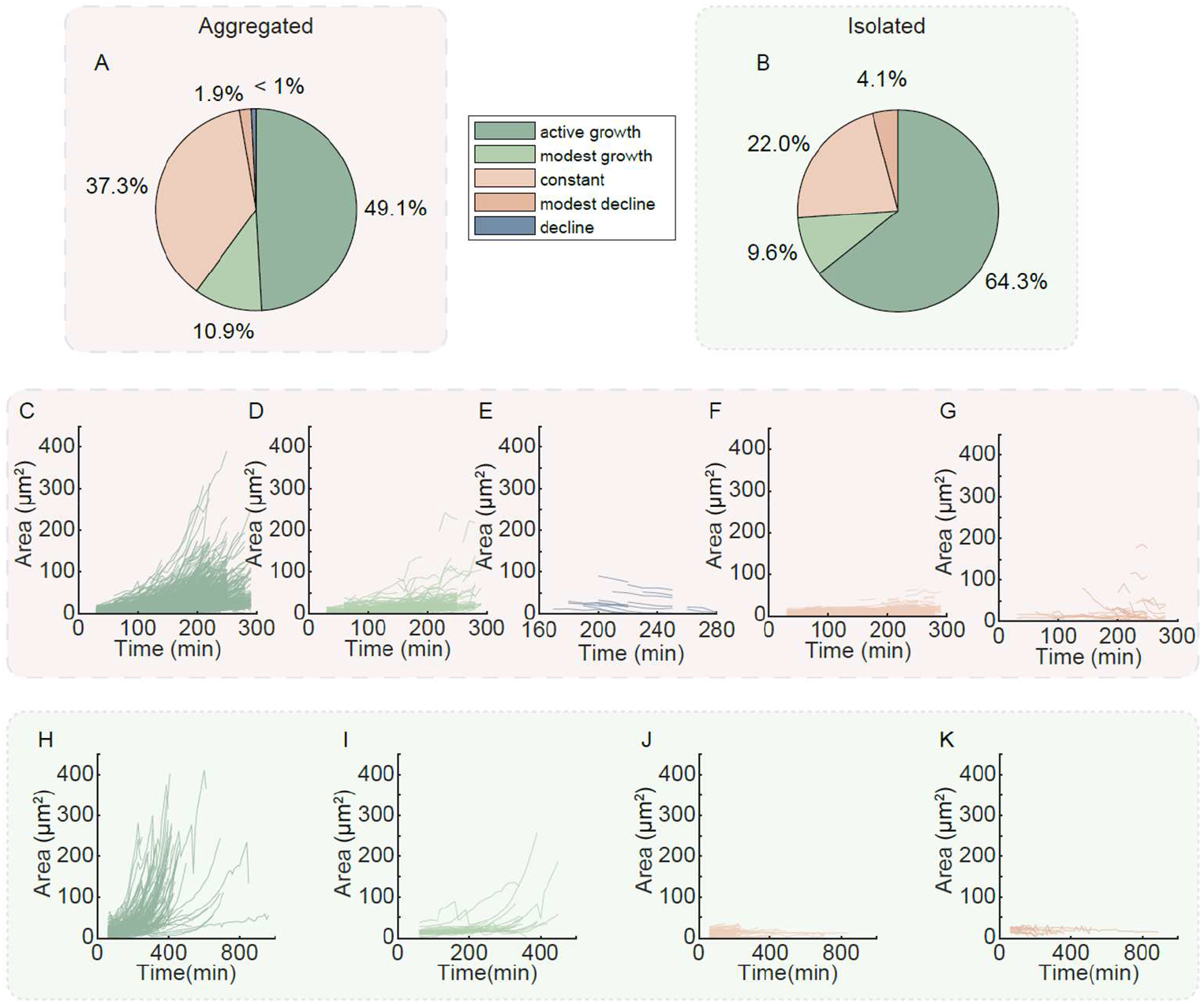
Analysis of growth dynamics of bacterial protocells in aggregates and in isolation. (A,B) Pie chart showing the proportions of five categories of protocell volume growth dynamics in aggregated protocolonies (panel A; N=1762) and in isolation (panel B; N=291). The classification criteria were based on the temporal dynamics of cell volume growth shown in panels C-K (Methods). The color coding in the legends applies to pie charts in both panels. (C-K) Temporal dynamics of cytoplasm-occupied area during the growth of cells in protocolonies (panels C-G) and in isolation (panels H-K). The color coding of the growth curves in panels C-K is the same as that in panels A,B. Each line corresponds to data from an individual cell, with T=0 min corresponding to the time of inoculation. The cytoplasm-occupied area, which is proportional to cell volume given the quasi-2D geometry, was measured by cell profiling based on fluorescence images of the cytoplasm stain CellROX Deep Red (Methods). Characteristics of each growth curve were evaluated, including standard deviation of data in the time sequence, growth rate obtained by exponential fit of the growth curve, and goodness of growth rate fitting (Methods). Cells were then classified into five categories according to criteria set based on these characteristics (Methods), which form the basis of color coding in all panels of this figure.

**Fig. S4.**
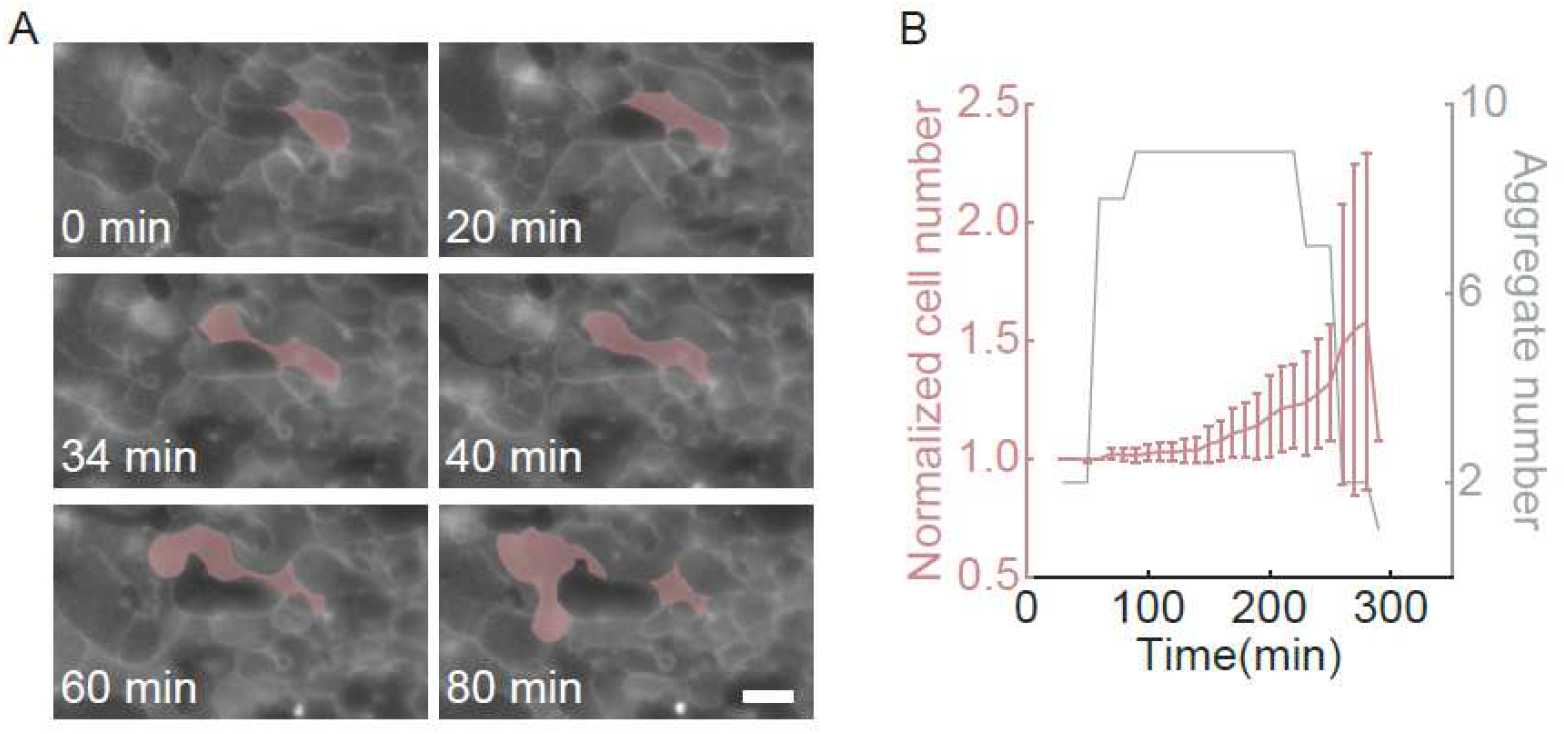
Division of cells in a protocolony. (A) Fluorescence image sequence showing the morphological change of CellROX-stained bacterial protocells in a protocolony (Methods). As shown in the images, CellROX fluorescence intensity is higher at the interface between cells, which allows for efficient cell segmentation. A representative protocell undergoing division is highlighted in red. T=0 min corresponds to the start time of the recording. Scale bar, 5 μm. (B) Cell number in protocolonies plotted against time. T=0 min corresponds to the time of inoculating the protocolonies. For each protocolony, the cell number was normalized by the cell number at the beginning of the observation. The data in red color (associated with left vertical axis) plots the average normalized cell number of protocolonies over time; error bars represent standard deviation (N>50 protocolonies), and lines connecting the data points serve as guides to the eyes. Data were acquired in 9 protocolonies of variable duration of observation, and the gray line (associated with the right vertical axis) plots the number of protocolonies from which the data point at a specific time was acquired. The result in panel B shows that the cell number in protocolonies started to increase at ∼2.5 hr after inoculation, suggesting that cell division events took place primarily after ∼2.5 hr post inoculation.

**Fig. S5.**
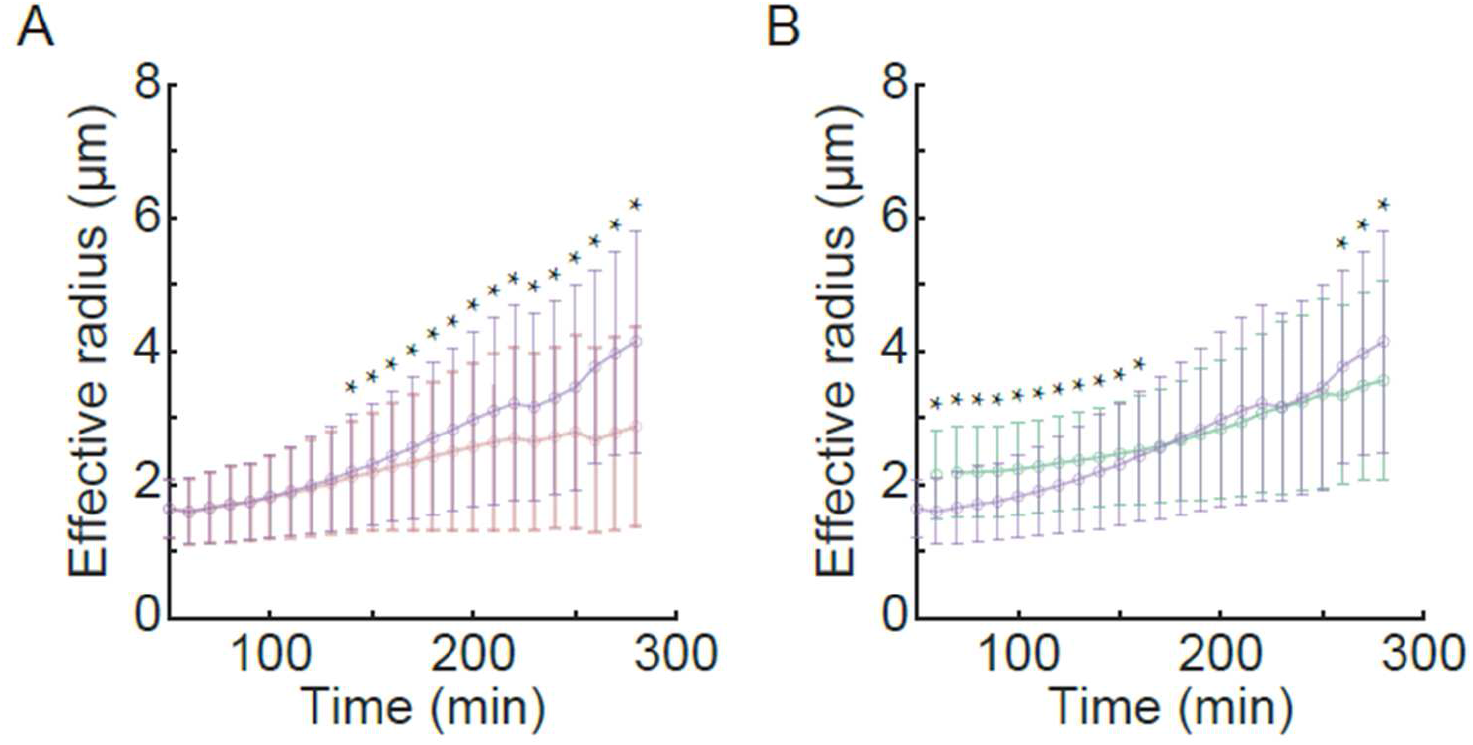
Impact of cell division on cell size in protocolonies. In both panels T=0 min corresponds to the time of inoculation, error bars represent standard deviation (N>50 cells), and lines connecting the data points serve as guides to the eyes. (A) Effective radius of cells in protocolonies plotted against growth time. The size or effective radius of a bacterial protocell is defined as 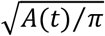, where *A*(*t*) is 2D projected cell area (Methods). Data in red color plot the actual average cell size, and data in violet color plot the average effective radius of virtual mother cells calculated based on the total area of daughter cells belonging to the same lineage (i.e., the area of a virtual mother cell at a specific time is the sum of all its surviving off-springs). Significance test for each pair of red and violet data sets at a specific time point was performed by two-tailed Mann-Whitney-Wilcoxon test to examine whether the two data sets were drawn from the same distribution (∗ *P* < 0.05). As shown in this panel, the red and violet data sets started to deviate from each other at T= ∼140 min, which corresponds to the onset of division events in the densely packed population; this result is consistent with data shown in Fig. S4B. (B) Comparison of the protocell size in aggregates and in isolation. Data in violet and green color correspond to virtual mother cells in protocolonies and isolated cells, respectively; the violet-color data are the same as those shown in panel A. Note that data of virtual mother cells are used for the comparison here in order to eliminate the impact of cell division on the cell size distribution. Significance test for each pair of green and violet data sets at a specific time point was performed in the same manner as in panel A. This panel shows that the average size of virtual mother cells in protocolonies is significantly smaller than that of isolated protocells during the initial growth stage (from T= 0 min to ∼150 min), but then it became comparable or even greater after T=∼150 min. Combining both panels of this figure and the main text Fig. 3A, we conclude that cell division is the primary cause of the significant size difference between cells in protocolonies and isolated protocells at the later stage of growth (> ∼150 min); the initial cell size difference (before ∼150 min) is presumably due to the fact that a larger proportion of cells in protocolonies were non-growing (37.3% in protocolonies vs. 22.0% for isolated cells; see Fig. S3A,B).

### Supplementary Table

**Table S1.** Parameters for Cellular Potts Model simulations.

| General parameters | Symbol | Value |  |  |  |
| --- | --- | --- | --- | --- | --- |
| System size (lattices) |  | 500×500 |  |  |  |
| Number of attempts per Monte Carlo Step (MCS) | | 8 ($\times 500^2$) | | | |
| Mean of initial cell radius |  | 5 |  |  |  |
| S.T.D. of initial cell radius |  | 1 |  |  |  |
| S.T.D. of initial distance between cells (center to center) |  | 0.5 |  |  |  |
| Temperature | $T$ | 2 | | | |
| Cell-environment contact energy | $J_{cell\&env}$ | 1 | | | |
| Cell-cell contact energy | $J_{cell\&cell}$ | 2 | | | |
| Buffering area in the area free energy $\mathcal{H}_{Area}$ | $A_B$ | 1.00E-07 | | | |
| Weight of volume growth | $\lambda_A$ | $\frac{4}{5\pi}$ | | | |
| Weight of perimeter growth | $\lambda_P$ | $\frac{2}{\pi}$ | | | |
| Coefficient of substrate-pinning energy | $\lambda_L$ | 2 | | | |
| Decay of weight of substrate-pinning energy per MCS | $\mu$ | 0.1 | | | |
| Increase of weight of substrate-pinning energy per MCS | $\gamma$ | 0.4 | | | |
| Time duration of each MCS | $\Delta T$ | $\frac{1}{12}$ | | | |
| Parameters for specific simulations | Symbol | Value or range |  |  |  |
|  |  | Fig. 5B | Fig.5C,E | Fig. 5D | Fig. 5F |
| Seeding cell number |  | 120 | 60 | 120 | 20 |
| Volume growth rate | $\beta_V$ | 0.6 | [0.1-0.8] | 0.6 | 0.6 |
| Mean of initial distance between cells (center to center) | $D_{cell}$ | 10.5 | 10.5 | 10.5 | [12-18] |
| S.T.D. of volume growth rate | $\beta_{V,sigma}$ | 0.1 | 0.001 | 0.1 | 0.1 |
| MCS steps |  | 30 | 60 | 60 | 60 |

### Supplementary Movies

**Movie S1. Growth dynamic of cells in a representative protocolony**. This video combines images acquired simultaneously from 3 channels: phase contrast (left), fluorescence of nucleic acids (middle; stained by SYTO16; Methods), and fluorescence of cell cytoplasm (right; stained by CellROX Deep Red; Methods). The video is played at 5 frames per second with the real elapsed time indicated in the time stamp. It is associated with Fig. 1A (upper) in main text. Scale bar, 50 μm.

**Movie S2. Growth dynamic of representative isolated bacterial protocells**. This video combines images acquired simultaneously from 3 channels: phase contrast (left), fluorescence of nucleic acids (middle; stained by SYTO16; Methods), and fluorescence of cell cytoplasm (right; stained by CellROX Deep Red; Methods). The video is played at 5 frames per second with the real elapsed time indicated in the time stamp. It is associated with Fig. 1A (lower) in main text. Scale bar, 20 μm.

